# Functional histamine H_3_ and adenosine A_2A_ receptor heteromers in recombinant cells and rat striatum

**DOI:** 10.1101/171736

**Authors:** Ricardo Márquez-Gómez, Citlaly Gutiérrez-Rodelo, Meridith T. Robins, Juan-Manuel Arias, Jesús-Alberto Olivares-Reyes, Richard M. van Rijn, José-Antonio Arias-Montaño

## Abstract

In the striatum, histamine H_3_ receptors (H_3_Rs) are co-expressed with adenosine A_2A_ receptors (A_2A_Rs) in the cortico-striatal glutamatergic afferents and the GABAergic medium-sized spiny neurons that originate the indirect pathway of the basal ganglia. This location allows H_3_Rs and A_2A_Rs to regulate the striatal GABAergic and glutamatergic transmission. However, whether these receptors interact to modulate the intra-striatal synaptic transmission has not yet been assessed. To test this hypothesis a heteromer-selective *in vitro* assay was used to detect functional complementation between a chimeric A_2A_R_302_-Gα_qi4_ and wild-type H_3_Rs in transfected HEK-293 cells. H_3_R activation with the agonist RAMH resulted in Ca^2+^ mobilization (pEC_50_ 7.31 ± 0.23; maximal stimulation, Emax 449 ± 25 % of basal) indicative of receptor heterodimerization. This response was not observed with histamine, suggesting a RAMH bias for heteromers. Functional A_2A_R-H_3_R heteromers were confirmed by co-immunoprecipitation and observations of differential cAMP signaling when both receptors were co-expressed in the same cell. In membranes from rat striatal synaptosomes, H_3_R activation decreased A_2A_R affinity for the agonist CGS-21680 (pKi values 8.10 ± 0.04 and 7.70 ± 0.04). Moreover, H_3_Rs and A_2A_Rs co-immunoprecipitated in protein extracts from striatal synaptosomes. These results support the existence of a H_3_R/A_2A_R heteromer, and reveal a new mechanism by which these receptors may modulate the unction of the striatum and the basal ganglia.

## 1. Introduction

The actions of histamine in the periphery on airway constriction, inflammation and gastric acid secretion are well known, whereas its function in the Central Nervous System (CNS) is not yet fully understood. In mammals, histamine actions are mediated by four (H_1_R-H_4_R) G protein-coupled receptors (GPCRs), of which the H_3_ receptor (H_3_R) displays the highest expression in the brain. In the CNS histamine is released from the terminals and dendrites of neurons located in the hypothalamic tuberomammilary nucleus, from where they innervate most of the CNS, including nuclei belonging to the basal ganglia, a subcortical group of structures intimately related to the control of motor behavior (Panula and Nuutinen, 2013).

The striatum is the main input nucleus of the basal ganglia and integrates motor and sensory information. The primary striatal afferents are glutamatergic axons originating from neurons located in the cerebral cortex and thalamus, and dopaminergic axons of sustantia nigra pars compacta neurons (Bolam *et al*., 2000). In turn, the axons of the GABAergic medium-sized spiny neurons (MSNs), which account for more than 90% of the striatal neuronal cell population (Kemp and Powell, 1971), project to the globus pallidus and sustantia nigra pars reticulata (Bolam *et al*., 2000).

H_3_Rs are abundantly expressed in the striatum either as pre-synaptic auto- and heteroreceptors or as post-synaptic receptors, with the latter located on the bodies of the MSNs and GABAergic or cholinergic interneurons (Pillot *et al*., 2002; González-Sepúlveda *et al*., 2013; Bolam and Ellender, 2016). The understanding of the physiological role of striatal H_3_Rs is complicated by their ubiquitous expression and capability to engage in protein-protein interactions with other GPCRs (Nieto-Alamilla *et al*., 2016). Of particular interest in the striatum are dopamine and adenosine receptors, which are also highly expressed. Based on the selective expression of dopamine and adenosine receptor subtypes, MSNs can be segregated into two neuronal populations. Neurons that originate the basal ganglia direct pathway (dMSNs) are identified by the expression of D_1_ receptors (D_1_Rs) whereas the indirect pathway population (iMSNs) is formed by neurons expressing adenosine A_2A_ receptors (A_2A_Rs) and D_2_ receptors (D_2_Rs) (Schiffman *et al*., 1991; Ferre *et al*., 1997; Tapper *et al*., 2004). In the striatum H_3_Rs have been shown to form heterodimeric protein-protein complexes with dopamine D_2_-like and D_1_-like receptors (Ellenbroek, 2013), and heterotrimers with σ1/D_1_ receptors and NMDA/D_1_ receptors (Moreno *et al*., 2014; Rodríguez-Ruíz *et al*., 2016). In turn, A_2A_Rs can form dimers with D_2_, cannabinoid CB_1_ and adenosine A_1_ receptors (Ciruela *et al*., 2006; Carriba *et al*., 2007; Hillon *et al*., 2002) and oligomers with D_2_ and mGlu5 glutamate receptors (Cabello *et al*., 2009).

Both H_3_ and A_2A_ receptors modulate intra-striatal synaptic transmission. The activation of Gα_i/o_-coupled H_3_Rs results in inhibition of GABA, dopamine, acetylcholine and glutamate release (Doreulee *et al*., 2001; Schlicker *et al*., 1993; Prast *et al*., 1999; Arias-Montaño *et al*., 2001), whereas the activation of Gα_olf_-coupled A_2A_Rs also inhibits GABA release but facilitates glutamatergic cortico-striatal transmission (Kirk and Richardson, 1994; Popoli, 1995).

Either individually or through their heteromeric interactions with other GPCRs, A_2A_Rs and H_3_Rs have been proposed as targets for the treatment of basal ganglia-related disorders such as Parkinson’s disease and addiction (Ferré *et al*., 2004; Schwarzschild *et al*., 2006; Passani and Blandina, 2011; Ellenbroek, 2013). In this work, we hypothesized that striatal H_3_Rs and A_2A_Rs can not only form heteromers with dopamine receptors, but also with each other, and that this interaction forms a unique pharmacological target that could be of potential therapeutic interest. Here we show that H_3_Rs co-immunoprecipitate with A_2A_Rs in HEK293 cells, suggestive of oligomerization. Further, we studied the physical properties of the A_2A_R/H_3_R interaction by functional complementation assay, in which we found an intriguing example of biased agonism. We next show that the A_2A_R/H_3_R heteromerization has unique functional pharmacology in terms of cAMP modulation in HEK-293 cells coexpressing the receptors. Using protein extracts from striatal synaptosomes, we confirmed the presence of A_2A_R/H_3_R heteromers *in vivo* by co-immunoprecipitation. Finally, we found that in the same preparation H_3_R activation resulted in decreased binding affinity of A_2A_R for the agonist CGS-21680.

A preliminary account of this work was presented in the abstract form to the European Histamine Research Society (Márquez-Gómez et al., 2016).

## 2. Methods

### 2.1 Materials

The following drugs and reagents were purchased from Sigma Aldrich (St. Louis, MO, USA): (R)-α-methyl-histamine dihydrochloride, histamine dihydrochloride, adenosine deaminase (from bovine spleen), Percoll, quinpirole dihydrochloride and caffeine. Dimaprit was from Axon MedChem (Reston, VA, USA). CGS-21680 was from Cayman Chemical (Ann Arbor, MI, USA). N-α-[methyl-^3^H]-histamine (78.3 Ci·mmol^−1^) and CGS-21680-[carboxyethyl-^3^H (N)]-(35.2 Ci·mmol^−1^) were from Perkin Elmer (Boston, MA, USA).

### 2.2 Molecular cloning

Truncated A_2A_ or H_3_ receptors were generated by PCR using the primers indicated in Supplementary Table 1. After amplification and restriction with HindIII and BamHI enzymes, the truncated A_2A_ and H_3_ receptors were ligated to a pcDNA3.1 plasmid that contained the chimeric Gα_qs4_ and Gα_qi4_ proteins. Both A_2A_Rs and H_3_Rs (cDNA Resource Center, Bloomsberg, PA, USA) were labeled with a triple hemagglutinin tag (3xHA). To generate the 3xHA tagged receptors, A_2A_R and the H_3_R were amplified from nucleotide 377 and 458, respectively, towards their amino terminus. The 3xHA tag was amplified from a plasmid that codified for the tagged histamine H_4_R (3xHA-H_4_R in pcDNA3.1, cDNA Resource Center). The amplified DNA fragments (3xHA-A_2A_R377/H_3_R458) where then reintroduced into the pcDNA3.1-H_3_R or pcDNA3.1-A_2A_R backbone to obtain the tagged, full length receptors. The insertion and orientation of the amplified fragment was verified by automated sequencing performed at FESI-UNAM (Los Reyes Iztacala, Estado de México, México).

**Table 1.** Pharmacological characteristics of the A_2A_R/H_3_R functional complementation assays in transfected HEK-293 cells

| Transfection | Agonist | E<sub>max</sub> (%) | pEC<sub>50</sub> |
| --- | --- | --- | --- |
| H<sub>3</sub>R + Gα<sub>qi4</sub> | RAMH | 419 ± 25 | 7.73 ± 0.31 |
|  | Histamine | 504 ± 22<sup>b</sup> | 6.19 ± 0.17<sup>b</sup> |
| H<sub>3</sub>R<sub>427</sub>-Gα<sub>qi4</sub> | RAMH | 753 ± 41 | 7.14 ± 0.21 |
| H<sub>3</sub>R<sub>427</sub>-Gα<sub>qi4</sub> + H<sub>3</sub>R | RAMH | 523 ± 41<sup>b</sup> | 7.38 ± 0.34 |
| A<sub>2A</sub>R<sub>302</sub>-Gα<sub>qi4</sub> + H<sub>3</sub>R | RAMH | 449 ± 25 | 7.31 ± 0.23 |
|  | RAMH + CGS | 429 ± 14<sup>n.s.</sup> | 7.63 ± 0.15 |
| H<sub>4</sub>R + Gα<sub>qi4</sub> | Histamine | 725 ± 51 | 6.35 ± 0.25 |
|  | RAMH | 551 ± 27<sup>a</sup> | 6.15 ± 0.16 |
| H<sub>2</sub>R + Gα<sub>qs4</sub> | Dimaprit | 860 ± 37 | 6.15 ± 0.12 |
| A<sub>2A</sub>R<sub>302</sub>-Gα<sub>qi4</sub> + D<sub>2</sub>R | Quinpirole | 612 ± 34 | 6.47 ± 0.22 |

Data are means ± SEM from 3 experiments. Statistical comparisons were performed between two agonists for the same transfection, or between the transfection with the chimeric receptor and the transfection of the chimeric receptor plus the wild type receptor.
<sup>a</sup> $P < 0.05$ ; <sup>b</sup> $P < 0.01$ , Student's *t* test. Both RAMH and CGS-21680 were assayed at 100 nM. RAMH, R-α-methylhistamine; CGS, CGS-21680.

### 2.3 Cell culture and transfection

HEK-293T cells (American Type Culture Collection, Manassas, VA, USA) were grown in Dulbecco’s modified Eagle’s medium (DMEM) supplemented with 10% fetal bovine serum (FBS), penicillin (50 UI/ml) and streptomycin (0.1 mg/ml) under a humidified atmosphere (5% CO_2_ in air) at 37 °C.

For transfections, cells were seeded in 6-well plates (6×10^5^ cells/well; up to passage 15) and incubated for 24 h at 37 °C in a 5% CO_2_/air mixture. The next day, 10 μl X-tremeGENE (Roche, Basilea, Switzerland) were mixed with 500 μl Optimem (Life Technologies, San Diego, CA, USA) and the mixture was incubated for 5 min at room temperature before the addition of cDNA in a 1:5 ratio with X-tremeGENE. For cAMP assays a mixture of DNA:Glosensor cAMP plasmid (1:2 ratio) was added to the Optimem/X-tremeGENE solution. When plasmids containing the A_2A_R or H_3_R were transfected alone, the amount of DNA was preserved by adding empty pcDNA3.1 vector. The transfection mixture was incubated for 20 min at room temperature and then added to the cells, which were incubated for 24 h at 37 °C under a humidified atmosphere (5% CO_2_/air).

For co-immunoprecipitation assays, HEK-293 cells, grown in 100-millimeter Petri dishes and at 80% confluence, were transfected using the polyethylenimine (PEI) method. Briefly, a mixture of DNA/PEI (1:10 ratio) was incubated for 30 min at room temperature before being added to the cells. After incubation for 30 min at 37 °C under a humidified atmosphere (5% CO_2_/air), cells were supplemented with 10% FBS and incubation continued for further 24 h under the same condition.

### 2.4 cAMP accumulation assay

cAMP assays were performed as described in detail elsewhere (Chiang et al. 2016). Briefly, HEK-293 cells transfected with 3xHA-A_2A_R, 3xHA-H_3_R or a mixture of both plasmids (1 μg each) together with Glosensor cAMP plasmid (Promega, Madison, WI, USA) were seeded in a white 384-well low-volume plate (25,000 cells/well in 7.5 μl medium) and incubated with Glo-equilibrium medium (7.5 μl, Promega). Drugs under test were added in a 5 μl volume (4x in HBSS solution) and endogenous cAMP luminescence was measured in real-time in a Flexstation 3 apparatus (Molecular Devices, Sunnyvale, CA, USA).

### 2.5 Calcium mobilization assay

HEK-293T cells transfected with 1 μg of DNA were seeded (2.5×10^4^ cells/well) in a 384-well clear bottom black plate (25 μl-volume) and incubated for 24 h at 37 °C in a humidified 5% CO_2_/air atmosphere, covered with Areaseal film (Sigma, St. Louis, MO, USA). Cells were then loaded with 25 μl of the FLIPR Ca^2+^ dye (Molecular Devices, Sunnyvale, CA, USA) and incubated for 1 h at 37 °C. Agonists were added in a 20 μl volume and calcium mobilization was measured in real-time for 2 min in a Flexstation 3 apparatus (Molecular Devices, Sunnyvale, CA, USA) and analyzed as described in detail in van Rijn *et al*. (2013).

### 2.6 Synaptosome preparation

Wistar rats (males, 250-300 g, provided by the Unidad de Productión y Experimentatión para Animales de Laboratorio; UPEAL-Cinvestav, Mexico City) were decapitated and the brain was quickly removed from the skull and the forebrain was separated and deposited on a metal plate placed on ice. The striata from 3-5 animals were dissected using forceps and the tissue was placed in 5 ml 0.32 M sucrose solution containing 10 mM Hepes, 1 mg/ml bovine serum albumin and 1 mM EDTA (pH 7.4 with NaOH). The tissue was homogenized using 10 strokes of a hand-held homogenizer (400 rpm), the homogenate was centrifuged (1000x*g*, 10 min, 4 °C) and the supernatant was pelleted at 14,000x*g*(12 min, 4 °C). The pellet was re-suspended in 5 ml of a Percoll solution (45 %, v:v), in Krebs-Henseleit-Ringer buffer (in mM: NaCl 140, Hepes 10, D-glucose 5, KCl 4.7, EDTA 1, pH 7.3 with NaOH). After centrifugation (2 min, 14,000x*g*, 4 °C), the upper phase was collected and brought up to 20 ml with Krebs-Ringer-Hepes (KRH) solution (in mM: NaCl 113, NaHCO_3_ 25, Hepes 20, D-glucose 15, KCl 4.7, CaCl_2_ 1.8, MgCl_2_ 1.2, KH_2_PO_4_ 1.2, pH 7.4 with NaOH). The suspension was centrifuged (20,000x*g*, 20 min, 4 °C) and the pellet (synaptosomes) was resuspended in KRH solution unless otherwise indicated.

### 2.7 Electron microscopy

Striatal synaptosomes were isolated by the Percoll method as above. Sample preparation and electron microscopy were performed as described in detail elsewhere (Morales-Figueroa *et al*., 2015).

### 2.8 Radioligand binding assays with striatal membranes

Striatal synaptosomes were re-suspended in lysis solution (Tris-HCl 10 mM, EGTA 1 mM, pH 7.4) and incubated for 20 min at 4 °C before centrifugation (20 min, 20,000xg, 4 °C). The pellet (synaptosomal membranes) was re-suspended in 1 ml KRH solution containing adenosine deaminase (2 U/ml). After incubation for 30 min at 37 °C, the suspension was brought to 20 ml with KRH solution and centrifuged (20 min, 20,000x*g*, 4 °C), the membranes were re-suspended in incubation solution (Tris-HCl 50 mM, MgCl_2_ 5 mM, pH 7.4) and 130 μl aliquots (300 μg protein) were incubated with 10 μl of increasing concentrations of CGS-21680 or RAMH (20x) and 50 μl of a fixed concentration of [^3^H]-NMHA (8 nM) or [^3^H]-CGS-21680 (48 nM). After 1 h at 30 °C ([^3^H]-NMHA) or 2 h at 25°C ([^3^H]-CGS-21680), incubations were stopped by rapid filtration through Whatman GF/B filters pre-soaked (2 h) in 0.3 % polyethylenimine (PEI). Filters were washed 3 times with 1 ml ice-cold buffer solution (50 mM Tris-HCl, pH 7.4), soaked in 4 ml scintillation solution and the tritium content was determined by scintillation counting.

### 2.9 Co-immunoprecipitation assays

Striatal synaptosomes were obtained as described above and then re-suspended in 1 ml of lysis solution (Tris-HCl 50 mM, NaCl 150 mM, Triton X-100 1%, SDS 0.05%, protease inhibitor 1 μl/ml). HEK-293 cells were dislodged in RIPA solution. For both synaptosomal and cell samples, protein was extracted by sonication (3 cycles, 30 sec, 8 kHz). The sample was centrifuged (20 min, 6,000x*g*) and the protein extract (supernatant) was collected. Nonspecific binding was removed by incubation (1 h, 4 °C) with 10 μl of AG beads (Santa Cruz Biotechnology; Dallas, TX, USA) under rotatory rocking. The beads were pelleted (2 min, 6000x*g*) and the supernatants were used for the assay. Protein quantification was performed by the BCA method.

The protein extracts (500 μg protein from striatal synaptosomes and 200 μg from HEK-293) were incubated with 2 μg of the primary antibodies (anti-A_2A_R abcam, cat. ab3461, lot. GR238882-9; anti-HA, Cell Signaling, cat. C29F4, lot. 3724S; anti-H_3_R, abcam, cat. ab84468, lot. GR27494-1, in a 1:100 dilution) together with AG beads (Santa Cruz Biotechnology sc-2003, 1:5 antibody to beads ratio) for 16 h at 4 °C with constant rotational rocking. Antibodies against the CD-81 protein (Santa Cruz Biotechnology, cat. sc-70803, lot. B0609) or D_2_R (Santa Cruz Biotechnology, cat. sc-5303, lot. A03013) were used as negative controls for the striatal synaptosomes or HEK-293 cells, respectively. The bead-antibody complex was pelleted (2 min, 6000x*g*) and a 30 μl aliquot of the supernatant was used as a load control. In both protein extracts, 30 μl of the total protein was used as input. The complex was dissociated for 60 min at 48°C in loading buffer (10% β-mercaptoethanol and 50% Laemmli buffer in H_2_O,) and separated by electrophoresis on a 10% SDS-polyacrylamide gel (20 min at 80 V and then 65 min at 120 V). Semi-dry transfer was performed at 15 V for 95 min. Membranes were blocked with 5% nonfat dry milk diluted in TBS-Twin 0.05% solution (overnight, 4 °C). After extensive washing, 5 ml of TBS-Twin 0.05% solution containing 5% BSA and the blotting antibody (anti-A_2A_R, anti-HA or anti-H_3_R, in a 1:1000 dilution) were added to the membrane and incubated overnight at 4 °C. Membranes were incubated with the secondary antibody (Invitrogen, HRP-conjugated anti-rabbit IgGs, cat. 65-6120, 1:5000, non-fat dry milk 5% in TBS-Twin 0.05%) at room temperature for 2 h. For the loading controls membranes were incubated for 1h with antibodies directed to β-tubulin (Invitrogen, cat. 322600, lot. 1235662) or α-actin (Sigma, cat. A5228, lot. 128K4843) in a 1:5000 dilution in TBST 0.05%. After washing, membranes were incubated for 1 h with the secondary antibody (α-mouse, Santa Cruz, cat. sc-2005) diluted in TBST 0.05%. Blot images were obtained by chemiluminescence in an X-ray film (Kodak).

### 2.10 Statistical analysis

All data was analyzed using the Prisma GraphPad software (San Diego, CA, USA). Data shows mean ± standard error of the mean (SEM). Statistical analysis was performed with student *t*-test.

## 3 Results

### 3.1 Co-immunoprecipitation of A_2A_Rs and H_3_Rs expressed in HEK-293 cells

In order to provide biochemical evidence for the hypothesized heteromerization of H_3_Rs and A_2A_Rs, we first studied by co-immunoprecipitation whether H_3_Rs and A_2A_Rs interact physically upon co-transfection in HEK-293 cells. The 3xHA-H_3_Rs were detected after immunoprecipitation of WT-A_2A_R (Figure 1A). No signal was observed when an antibody against the D_2_R (α-D_2_R) was employed as a negative control. These results support dimerization between H_3_Rs and A_2A_Rs.

**Figure 1.**
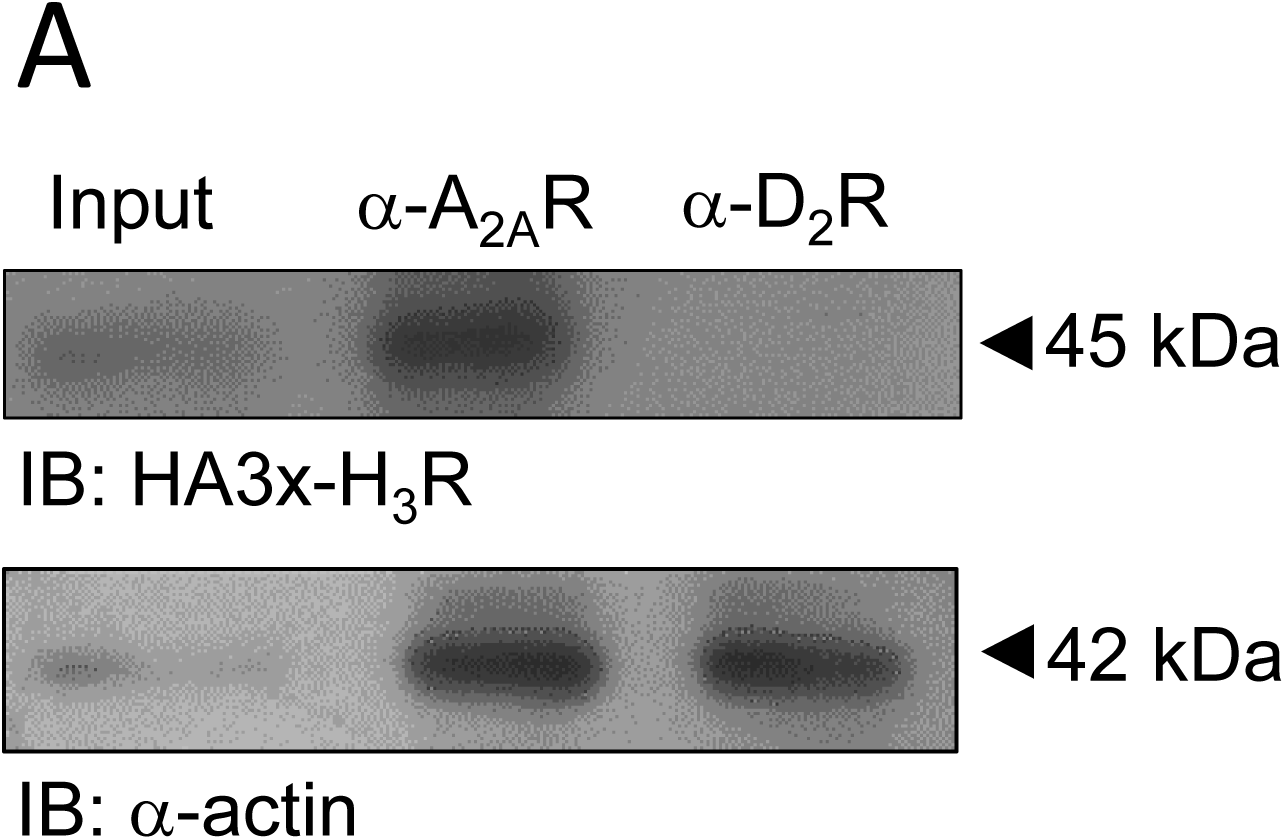
Adenosine A_2A_ and histamine H_3_ receptors co-immunoprecipitate in transfected HEK-293 cells. A. Co-immunoprecipitation of the hemagglutinin-tagged H_3_R (3xHA-H_3_R) with the A_2A_R in protein extracts from HEK-293 cells. A 45 kDa band corresponding to the A_2A_R was detected. This band was not observed with the negative control. Input corresponds to 30% of the total protein. An antibody against the D_2_ receptor (α-D_2_R) was used as a negative control. Blots are representative of 3 independent experiments.

### 3.2 Functional complementation of A_2A_Rs and H_3_Rs expressed in HEK-293 cells

To further study the potential interaction between A_2A_Rs and H_3_Rs in a recombinant cell system we employed a functional complementation assay, previously used to study physical interactions between GPCRs (Han *et al*., 2009; van Rijn *et al*., 2013). This assay relies on the fusion of chimeric Gα_qs4_ or Gα_qi4_ proteins to the truncated C-terminal tail of a GPCR to generate a nonfunctional receptor, rescuable through homo- or heterodimerization with a wild type (WT), untruncated receptor. It was previously reported that fusion of a chimeric G protein to a GPCR truncated near helix 8 produces a receptor that is unable to signal by itself but can be rescued when transfected with a full length receptor (van Rijn *et al*., 2013). Therefore, we first generated an A_2A_R truncated to 302 residues (A_2A_R_302_) and H_3_Rs of 427, 421 and 411 residues (H_3_R_427_, H_3_R_421_ and H_3_R_411_), and the truncated receptors were then fused to the chimeric G proteins (Figure 2A).

**Figure 2.**
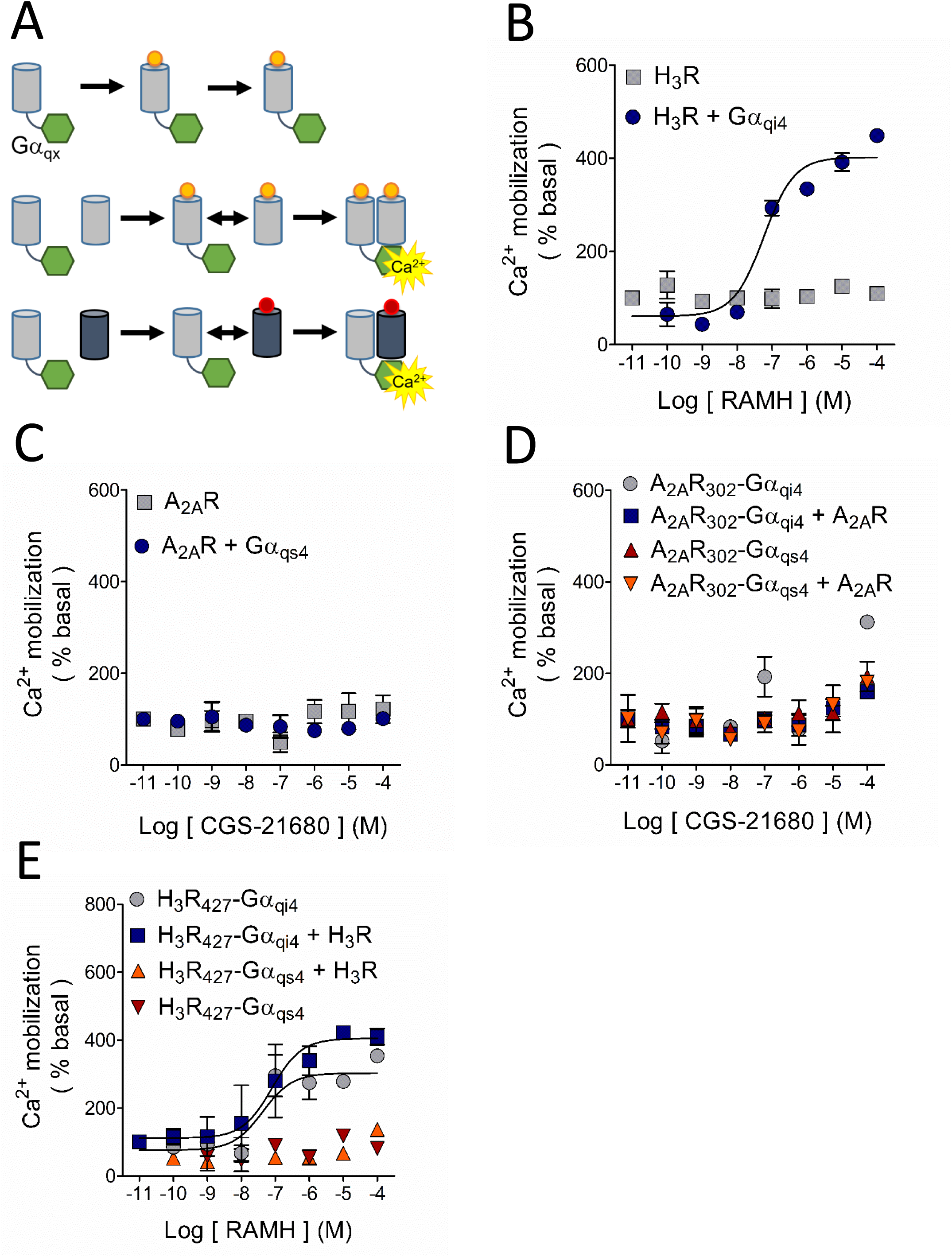
Study of the possible A_2A_R/H_3_R dimerization by functional complementation assay. A. Basis of the assay. The G protein-coupled receptor (GPCR; grey cylinders) is truncated at its carboxyl terminus to generate a nonfunctional GPCR, which is then fused to the chimeric G protein (green hexagons). The chimeric G protein is formed by a Gα_q_ protein in which 9 or 10 residues of the C-terminus were substituted by the corresponding sequence of a Gα_s4_ or Gα_i4_ protein (represented by X). Under these conditions, Ca^2+^ mobilization can only be elicited when the complex GPCR-Gα_qx_ is in close proximity to a wild type (WT) GPCR, either the same (homodimerization) or a different receptor (heterodimerization). B. Activation of the H_3_R with its agonist RAMH resulted in Ca^2+^ mobilization only when it was co-expressed with the chimeric Gα_qi4_ but not when transfected alone. C. When activated with its agonist CGS-21680, the A_2A_R was uncapable to induce Ca^2+^ mobilization either alone or when co-transfected with the chimeric Gα_qs4_ protein. D. Functional complementation by homodimerization of the truncated A_2A_R_302_, either bound to Gα_qs4_ (A_2A_R_302_-Gα_qs4_) or Gα_qi4_ (A_2A_R_302_-Gα_qi4_), with the native A_2A_R was not observed. E. Activation of the truncated H_3_R_427_ bound to Gα_qi4_ (H_3_R_427_-Gα_qi4_) induced Ca^2+^ mobilization on its own and homodimerization with the WT-H_3_R did not modify the response. No signal was observed with activation of the H_3_R_427_-Gα_qs4_ construct when it was co-expressed with the WT-H_3_R. In all graphs data are means ± SEM from 3 replicates from a representative experiment. The quantitative analysis is shown in Table 1 and 4.

Activation of the H_3_R in transfected CHO-K1 cells and rat striatal neurons in primary culture induces Ca^2+^ mobilization (Cogé *et al*., 2001; Rivera-Ramirez *et al*., 2016), and in the striatum A_2A_R activation favors Ca^2+^ entry by modulating voltage-activated Ca^2+^ channels (Kirk and Richardson, 1995; Gubitz *et al*., 1996), which are endogenously expressed by HEK-293 cells (Berjukow *et al*., 1996; Thomas and Smart, 2005). In HEK-293 cells the H_3_R selective agonist RAMH did not induced any discernible Ca^2+^ response but did so when the H_3_R was co-transfected with Gα_qi4_ proteins (Figure 2B). The A_2A_R did not induce Ca^2+^ signaling when activated by the selective agonist CGS-21680, but also no response was observed when co-transfected with Gα_qs4_ proteins, suggesting that A_2A_Rs do not activate this chimeric protein (Figure 2C). It has been reported that not all GPCRs are amenable to signal through chimeric G-proteins (Conklin *et al*., 1996). A different Gα_s_-coupled receptor, the histamine H_2_ receptor, was capable to induce calcium release when co-transfected with Gα_qs4_ proteins and stimulated with the selective agonist dimaprit (Supplementary Figure 1A), discarding Gα_qs4_ malfunction. Consistent with the finding that A_2A_Rs did not signal efficiently via chimeric G-proteins, functional complementation by homodimerization of A_2A_R_302_-Gα_qs4_ or A_2A_R_302_-Gα_qi4_ with the full length A_2A_R was not observed (Figure 2D).

The C-tail of the H_3_R was truncated such that the H_3_R-Gα_qi4_ fusion protein would display limited Ca^2+^ mobilization when expressed alone, but pronounced Ca^2+^ signaling when coexpressed with WT-H_3_Rs. The H_3_R_421_-Gα_qi4_ and the H_3_R_411_-Gα_qi4_ constructs did not induce Ca^2+^ mobilization, and could not be rescued via homo-dimerization with WT-H_3_Rs (Supplementary Figure 1B). However, Ca^2+^ mobilization induced by H_3_R_427_-Gα_qi4_ upon activation and co-transfection with WT-H_3_Rs produced a similar increase in calcium release (Figure 2E). While not optimal, this behavior allowed for the study of the interaction between the H_3_R_427_-Gα_qi4_ and the A_2A_R as explained below.

Co-transfection of the ‘inert’ A_2A_R_302_-Gα_qi4_ and H_3_Rs induced Ca^2+^ mobilization after activation of the latter receptors with RAMH, suggestive of close physical proximity between the H_3_R and both the A_2A_R and the fused Gα_qi4_-protein. H_3_R activation failed to induce functional complementation in cells co-transfected with A_2A_R_302_-Gα_qs4_ (Figure 3A). Given that H_3_Rs are Gα_i/o_-coupled, and the inability of the A_2A_R to signal through chimeric Gα_qi4_- or Gα_qs4_-proteins, it was not surprising that A_2A_R activation with the selective agonist CGS-21680 did not produce any response through H_3_R_427_-Gα_qs4_ or H_3_R_427_-Gα_qi4_ (Figure 3B). Interestingly, the co-expression of WT A_2A_Rs prevented RAMH-induced calcium mobilization mediated by the H_3_R_427_-Gα_qi4_ (compare Figure 2E and Figure 3C). We briefly explored this pharmacological response further by analyzing calcium release in HEK-293 cells co-expressing the H_3_R4_27_-Gα_qi4_ and the A_2A_R or the A_2A_R_302_-Gα_qi4_ and the H_3_R and activating both receptors, but did not observe any additive or antagonistic effect (Supplementary Figure 1C and 1D).

**Figure 3.**
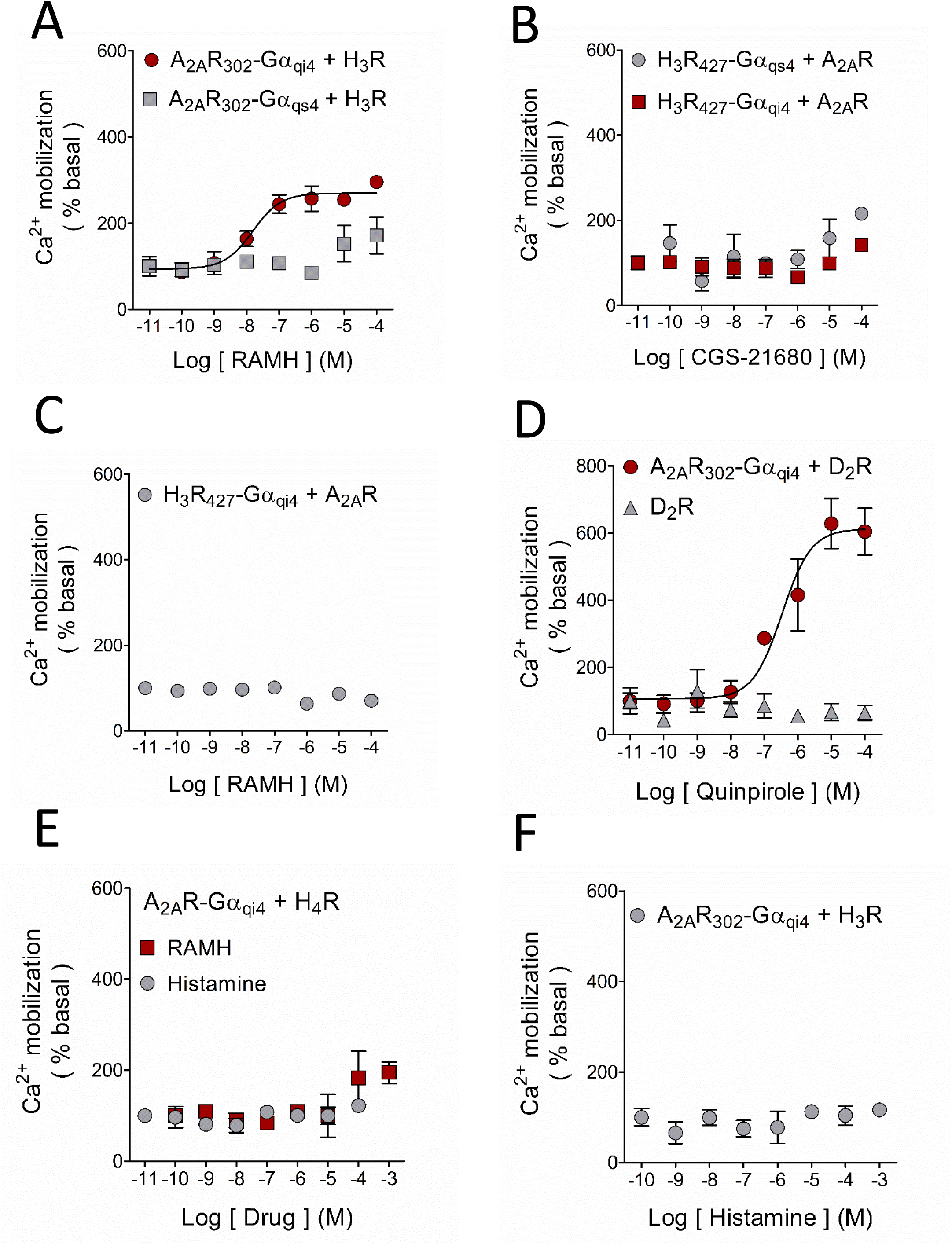
Ligand modulation of the A_2A_R/H_3_R interaction in HEK-293 cells. Activation of the WT-H_3_R with its agonist RAMH led to functional complementation when co-expressed with the A_2A_R_302_-Gα_qi4_ but not with A_2A_R_302_-Gα_qs4_. B. Ca^2+^ mobilization was not observed when the WT-A_2A_R was co-transfected with the truncated H_3_R_427_-Gα_qs4_ or H_3_R_427_-Gα_qi4_, in accord with the incapability of the receptor to activate the chimeric proteins. C. As shown in Figure 1D, activation of the H_3_R_427_-Gα_qi4_ resulted in Ca^2+^ mobilization, and this response was prevented when it was co-expressed with the WT-A_2A_R, suggesting a preference of the H_3_R_427_-Gα_qi4_ to form heterodimers. D. The well-studied A_2A_R/D_2_R heterodimer was used as a positive control for these experiments. Activation of the D_2_R with increasing concentrations of the agonist quinpirole caused marked Ca^2+^ mobilization only when coexpressed with the chimeric A_2A_R_302_-Gα_qi4_. E. H_4_R as a negative control. No functional complementation was observed when the receptor was co-expressed with the A_2A_R_302_-Gα_qi4_, showing the specificity of the A_2A_R/H_3_R interaction. F. Functional complementation between the A_2A_R_302_-Gα_qi4_ and the WT-H_3_R was not observed when the latter receptor was activated with the endogenous agonist histamine. For all graphs data are means ± SEM from 3 replicates from representative experiments. The quantitative analysis is shown in Table 1.

A_2A_Rs have been shown to form heteromers with D_2_Rs (Ferré *et al*., 2003) and robust functional complementation was accordingly observed when A_2A_R-Gα_qi4_ was co-expressed with WT-D_2_Rs (Figure 3D), supporting that our results are due to heterodimerization and not to stochastic interactions.

### 3.3 The putative A_2A_R/H_3_R heteromer displays ligand bias

To test for the selective nature of the A_2A_R/H_3_R interaction in the functional complementation assay, we investigated if A_2A_R-Gα_qi4_ would also allow the structurally and physiologically similar histamine H_4_ receptor (H_4_R) to induce calcium release. Coactivation with histamine of H_3_Rs and H_4_Rs elicited Ca^2+^ mobilization only when cotransfected with Gα_qi4_ (Supplementary Figure 2A and 2B, respectively). Neither histamine nor RAMH induced Ca^2+^ release when the H_4_R was co-expressed with A_2A_R_302_-Gα_qi4_, supporting the specificity of the A_2A_R/H_3_R interaction (Figure 3E). Unexpectedly, whereas RAMH induced Ca^2+^ mobilization when the H_3_R was co-expressed with A_2A_R_302_-Gα_qi4_, histamine did not (compare Figure 3A with Figure 3F). This result suggests that the A_2A_R/H_3_R heteromer displays agonist bias.

### 3.4 A_2A_R-mediated signaling is increased by H_3_R co-activation

To study the pharmacology of the putative A_2A_R/H_3_R heteromer we next tested cAMP signaling using WT receptors in HEK-293 cells. In cells transfected with A_2A_Rs, incubation with CGS-21680 resulted in a concentration-dependent increase in cAMP levels in accordance with the receptor’s coupling to the Gα_s_ signaling pathway (Figure 3A). CGS-21680 induced a similar cAMP response in HEK293-A_2A_R cells co-expressing H_3_Rs (Figure 4A). Co-activation of the receptors resulted in a decrease in baseline, in accordance with the Gα_i/o_ coupling of the H_3_R, but led to an augmentation of A_2A_R-mediated cAMP formation as noticed by a 2.5 fold change over baseline (versus 1.5 fold in control). This result suggests that the A_2A_R signaling becomes more efficacious when heterodimerization with the H_3_R occurs (Figure 4A, Supplementary Figure 2C, and Table 2). The increase in the A_2A_R signaling was not observed in the absence of RAMH (1.5 fold over baseline) suggesting an agonist-dependency of the response (Figure 4A and Supplementary Figure 2C).

**Figure 4.**
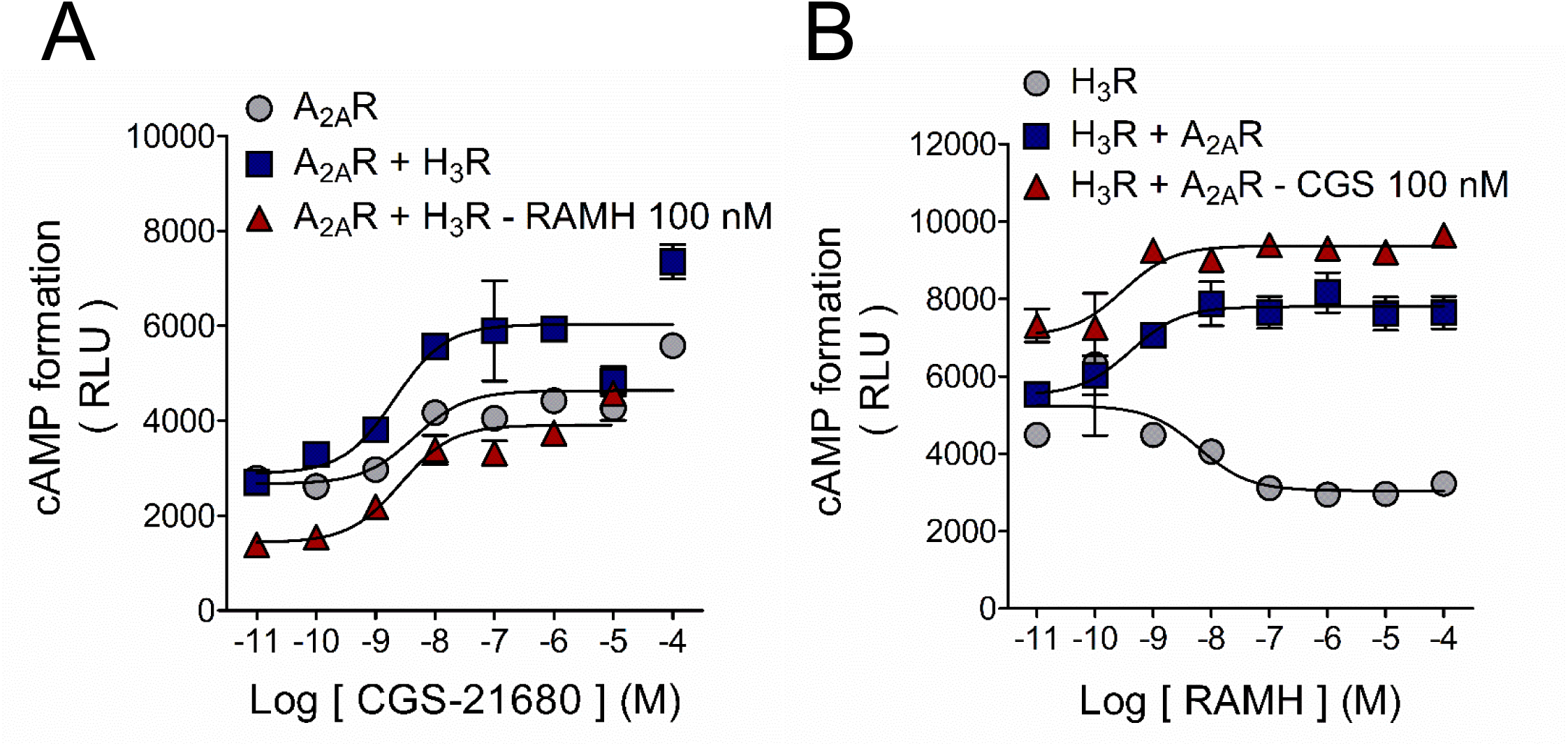
H_3_R activation enhances A_2A_R-mediated cAMP signaling in HEK-293 cells. A. Activation of the Gα_s_-coupled A_2A_R with its agonist CGS-21680 induced cAMP formation and H_3_R co-activation enhanced A_2A_R efficacy. Co-expression of the H_3_R did not modified the A_2A_R functional response. Values for pEC_50_ and maximal effect (Emax) are given in Table 2. B. H_3_R activation with RAMH decreased forskolin-induced cAMP formation in accord with the inhibitory nature of the Gα_i/o_-coupled receptor. A_2A_R expression and receptors co-activation lead to a change in the H_3_R signaling. Data are means ± SEM from 3-5 independent experiments. Values for pIC_50_ and maximal effect (Imax) are given in Table 2.

**Table 2.** Pharmacological characteristics of the cyclic AMP (cAMP) formation assay in transfected HEK-293 cells.

| Transfection | Agonist | pEC<sub>50</sub> | E<sub>max</sub> (%) | pIC<sub>50</sub> | I<sub>max</sub> (%) |
| --- | --- | --- | --- | --- | --- |
| A<sub>2A</sub>R | CGS | 8.0 ± 0.4 | 152 ± 7 | - | - |
| A<sub>2A</sub>R + H<sub>3</sub>R | CGS | 8.7 ± 0.5 | 153 ± 7 | - | - |
| A<sub>2A</sub>R + H<sub>3</sub>R | +RAMH | 8.0 ± 0.2 | 248 ± 9<sup>b</sup> | - | - |
| H<sub>3</sub>R | RAMH | - | - | 8.1 ± 0.3 | -26 ± 2 |
| H<sub>3</sub>R + A<sub>2A</sub>R | RAMH | 10.6 ± 1 | 114 ± 3 | - | - |
| H<sub>3</sub>R + A<sub>2A</sub>R | +CGS | 9.9 ± 0.6 | 120 ± 2 | - | - |

Data are means ± SEM from 3-5 experiments. Comparison was performed between the solely transfection of a receptor vs co-transfection of both receptors. N.D., not determined.
<sup>a</sup> $P < 0.05$ ; <sup>b</sup> $P < 0.01$ . Student's $t$ test. Both RAMH and CGS-21680 were tested at 100 nM.
RAMH, R- $\alpha$ -methylhistamine; CGS, CGS-21680; FSK, Forskolin.

In HEK293-H_3_R cells RAMH inhibited forskolin-stimulated cAMP formation congruent with the Gα_i/o_ coupling of the receptor. However, when the A_2A_R was co-transfected, RAMH-mediated H_3_R signaling shifted to facilitate cAMP formation instead. Activation of the A_2A_R caused an expected increase in the baseline but it did not significantly modify the cAMP-increase effect. This result suggests that in the heterodimer, A_2A_R signaling prevails over the H_3_R (Figure 4B; Table 2).

### 3.5 H_3_R activation decreases A_2A_R binding affinity in synaptosomal membranes

The identification of A_2A_R/H_3_R heteromers was performed in recombinant cell systems over-expressing the receptors. As previously mentioned A_2A_Rs and H_3_Rs are co-expressed in iMSNs and cortico-striatal projections. We therefore used rat striatal synaptosomes to seek for pharmacological traces that supported the existence of native A_2A_R/H_3_R dimers.

Electron microscopy confirmed in the purified synaptosomal preparation the vast presence and conserved structure of isolated nerve terminals, characterized by a delimited membrane and the presence of mitochondria and synaptic vesicles (Supplementary Figure 3A and B). In protein extracts from striatal synaptosomes immunoprecipitation of the A_2A_R resulted in a band of ~45 kDa, corresponding to the expected migration of the H_3_R. No signal was detected when an irrelevant antibody (α-CD81) was tested. As a positive control the H_3_R was immunoprecipitated and detected as a band of the same 45 kDa (Figure 5A). When the reverse approach was employed and the H_3_R was immunoprecipitated from the striatal protein extracts, a band of ~45 kDa corresponding to the expected migration of the A_2A_R was observed. This band was also observed when the A_2A_R was immunoprecipitated as a positive control but not detected in the negative control (Figure 5B). This result supports that the interaction A_2A_R/H_3_R is constitutively present in the striatal nerve terminals.

**Figure 5.**
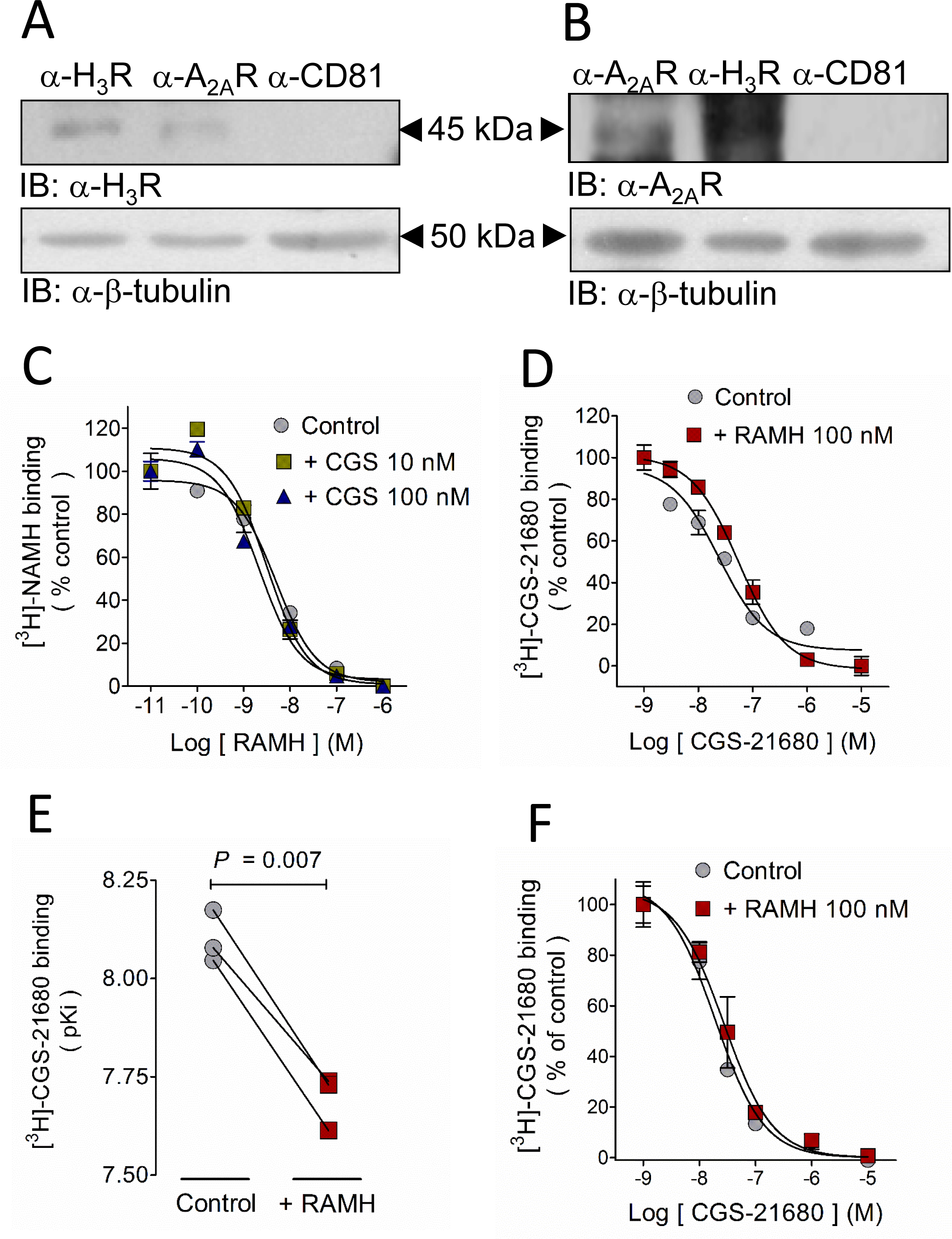
H_3_R activation decreases A_2A_R affinity for the agonist CGS-21680 in synaptosomal membranes but not in membranes from the whole striatum. A. The H_3_R co-immunoprecipitated with the A_2A_R in a protein extract of Percoll-purified striatal synaptosomes. The band of ~45 KDa corresponds to the expected migration of the H_3_R. No band was observed in the negative control (αCD81). B. Co-immunoprecipitation of the A_2A_R with the H_3_R in a protein extract of striatal synaptosomes. A band of ~45 KDa corresponds to the expected migration of the H_3_R.The figure depicts representative blots, repeated a further 4 times with similar results. C. CGS-21680 (10 and 100 nM) did not modify the H_3_R affinity for its ligand [^3^H]-NAMH in membranes isolated from striatal synaptosomes. D. In the same preparation, the H_3_R agonist RAMH (100 nM) decreased the A_2A_R affinity for [^3^H]-CGS-21680. Values are means ± SEM from 3 replicates from a representative experiment. E. Analysis of 3 independent experiments. The statistical analysis was performed with paired Student’s *t* test. F. H_3_R activation with RAMH (100 nM) failed to decreased the A_2A_R affinity for [^3^H]-CGS-21680 in membranes from the whole striatum. Values are means ± SEM from 3 replicates from a representative experiment.

In binding studies with synaptosomal membranes the A_2A_R agonist CGS-21680 (10 and 100 nM) did not affect the affinity of the H_3_R for its agonist RAMH (Figure 5C), whereas RAMH (100 nM) decreased by two-fold the affinity of the A_2A_R for CGS-21680 (Figure 5D, 5E and Table 3), with no effect on maximal binding (Bmax). This pharmacology appears to be specific for pre-synaptic H_3_Rs and A_2A_Rs, because we did not observe a similar decrease in A_2A_R affinity for CGS-21680 in membranes from the whole striatum, in which post-synaptic membranes constitute the major component (Figure 5F and Table 3).

**Table 3.** Analysis of binding assays in rat striatal membranes

|  | pKi | Bmax (%) |
| --- | --- | --- |
| **[<sup>3</sup>H]-NAMH** |  |  |
| Synaptosomes |  |  |
| RAMH | 9.09 ± 0.22 | 100.0 ± 0.3 |
| RAMH + CGS 10 nM | 8.99 ± 0.02<sup>ns</sup> | 110.0 ± 4.0<sup>ns</sup> |
| RAMH + CGS 100 nM | 9.08 ± 0.06<sup>ns</sup> | 103.0 ± 4.1<sup>ns</sup> |
| **[<sup>3</sup>H]CGS-21680** |  |  |
| Whole striatum |  |  |
| CGS | 8.06 ± 0.15 | 100.0 ± 14 |
| CGS + RAMH | 8.11 ± 0.04<sup>ns</sup> | 88.0 ± 6.0<sup>ns</sup> |
| Synaptosomes |  |  |
| CGS | 8.10 ± 0.04 | 100.0 ± 9.0 |
| CGS + RAMH | 7.70 ± 0.04<sup>a</sup> | 88.1 ± 10.0<sup>ns</sup> |

Data are means ± SEM from 3-5 experiments. For [<sup>3</sup>H]-NAMH binding assays the statistical analysis was performed with one-way Anova and Dunnett's *post hoc* test. For [<sup>3</sup>H]-CGS-21680 binding, values in the presence of RAMH were compared with the corresponding control with Student's *t* test. <sup>a</sup> *P* < 0.001, ns, not significant. CGS, CGS-21680. RAMH (R-α-methylhistamine) was tested at 100 nM in all the experiments.

## 4 Discussion

The H_3_R has been proposed as a potential novel drug target for the treatment of drug use disorders, depression, schizophrenia and Parkinson’s disease. However, its wide brain expression and effects on other neurotransmitter systems may result in adverse effects when H_3_R selective drugs are administered systemically. The A_2A_R has also been proposed as an option for the treatment of Parkinson´s disease and possesses a strategic distribution in the striatum for targeting and modulating the cortico-dMSNs projections and the activity of iMSNs. The results presented herein may lead to an alternative to specifically target H_3_Rs located in either iMSNs or cortical afferents synapsing onto dMSNs projections.

### 4.1 A_2A_R/H_3_R interaction

The primary finding of this study was the identification, for the first time, of A_2A_R/H_3_R heteromers, not only in recombinant cell systems but also in rat striatal nerve terminals.

The basis of the functional complementation assays requires for a receptor to be nonfunctional and this was obtained by truncation of the carboxyl terminus (CT). Herein we showed that truncation of the H_3_R from 445 to 411 or 421 residues was sufficient to prevent H_3_R-mediated G protein activation of the H_3_R-G_qi4_ fusion protein, yet we were unable to functionally rescue signaling, indicating that the remaining C-tail may be too short to connect with an interacting GPCR. However, truncation to 427 amino acids allowed the H_3_R to remain functional. It may be possible to create a truncated version of the H_3_R somewhere between amino acids 421-427 short enough to abolish homo/monomeric signaling but long enough to functionally complement with a full length GPCR. A requirement for helix 8 has been demonstrated for D_1_Rs and opioid κ and μ receptors (van Rijn *et al*. 2013). The H_3_R third intracellular loop appears to play a key role in the receptor-G protein coupling on the basis of the decreased signaling of the H_3_R_365_ isoform, which lacks 80 residues in the third intracellular loop (Riddy *et al*. 2016), and reduced signaling was induced by the A280V mutation in the same loop (Flores-Clemente *et al*. 2013). Further consideration to the H_3_R carboxyl tail should therefore be made when assessing receptor-G protein coupling.

Ca^2+^ mobilization was detected upon exposure to the H_3_R agonist RAMH in HEK-293 cells transfected with H_3_R_427_-Gα_qi4_ (Figure 1E). Whereas this may appear to be an undesirable result, it further supports the formation of A_2A_R/H_3_R heterodimers because H_3_R4_27_-Gα_qi4_ function was not observed after co-transfection with A_2A_Rs, suggesting a preference of the H_3_R to form hetero-over homo-dimers.

An important part of describing a new heterodimer is to find changes in the signaling profiles of the receptors involved in the dimer. In this regard, we found enhanced signaling efficacy of the agonist CGS-21680 when the H_3_R was co-transfected and co-activated in the HEK293-A_2A_R cells. This effect can be explained by a H_3_R-mediated facilitation of the A_2A_R coupling to G proteins, leading to an increase in the receptor efficacy to activate Gα_s_ proteins and thus to produce cAMP. Given that in the absence of RAMH, CGS-21680 behaved the same in HEK293-A_2A_R cells and in HEK293-A_2A_R/H_3_R cells, receptor expression does not appear to account for the observed effects.

Canonically H_3_R couples to Gα_i/o_ proteins which inhibit adenylyl cyclase activity and accordingly, activation of the receptor with RAMH lead to a decrease in cAMP formation. Similar to the observed in the calcium mobilization assays, expression of the A_2A_R changed the H_3_R-mediated cAMP response from inhibition to an increase in the cAMP formation. In the calcium assays we did not observed signaling of the H_3_R through the Gα_qs_4 proteins bound to the truncated A_2A_R, and we are thus not considering a H_3_R change in signaling pathways as an explanation for this effect. Therefore, we hypothesized that in the A_2A_R/H_3_R heteromer the A_2A_R signaling prevails over H_3_R, while the latter signaling is hampered possibly by a steric impediment. This is in line with the loss of calcium signaling when H_3_R-Gα_qi4_ was co-expressed with A_2A_R.

An unexpected but interesting finding was the potential biased-signaling at the A_2A_R/H_3_R heteromer, as evidenced by the incapability of the endogenous ligand histamine to signal at the A_2A_R/H_3_R heteromer compared to the exogenous agonist RAMH. This discrepancy suggests that RAMH induces conformational changes in the H_3_R that allow the interaction to occur, whereas the histamine-induced changes appear not sufficient for heteromerization or at least for the heteromer to signal. Ligand bias could also be explained by agonist residence time, this is, the time that a particular drug remains in its binding pocket and that will directly determine the time that a given receptor maintains an active conformation. The active state induced by histamine may therefore have a shorter duration compared with that induced by RAMH, preventing the former from activating the chimeric G proteins bound to the A_2A_R_302_. Binding studies with striatal membranes and histamine show that the first hypothesis holds better because histamine induced an increase in the affinity of the A_2A_R for its agonist CGS-21680 (Supplementary figure 4), opposite to the decrease in affinity change induced by RAMH in the same preparation. This result suggests that histamine and RAMH lock the H_3_R in different conformational states that affect its interaction with the A_2A_R.

### 4.2 Synaptic location of the A_2A_R/H_3_R interaction

A previous report indicates equal distribution of A_2A_Rs in total and synaptosomal membranes from rat striatum (Rebola *et al*., 2005), with a preferential location on the postsynaptic density fraction over the pre-synaptic active zone fraction (49.2 ± 3.3 % and 26.9 ± 3.3 % of total immunoreactivity, respectively). H_3_Rs are expressed pre- and post-synaptically (Ellenbroek and Ghiabi, 2014), but they seem to be highly enriched in the terminals of striato-pallidal neurons (iMSNs) yielding a value of 1,327 ± 79 fmol/mg protein (Morales-Figueroa *et al*., 2014). Given this distribution, our results point to a specific synaptic location of the A_2A_R/H_3_R heteromer that provides the interaction a preferential role in pre-synaptic modulation of glutamatergic or GABAergic transmission.

In the striatum, A_2A_Rs have a specific location in iMSNs (Schiffman *et al*., 1991) and cortico-dMSNs terminals, but not in the cortico-iMSNs projections (Quiróz *et al*., 2009). It has also been reported that the A_2A_R antagonists KW-6002 and SCH-442416 can differentiate either A_2A_R location, respectively (Orrú *et al*., 2011). Although there are no agonists capable to differentiate between A_2A_R populations, this information may be a valuable tool for targeting and modulating the A_2A_R and H_3_R pharmacology in a location-specific manner, using bivalent ligands directed to the heterodimer. Speculating on the potential therapeutic role of the A_2A_R/H_3_R -heteromer and assuming an increase in the A_2A_R signaling, targeting the cortico-striatal heteromer may have relevance for the treatment of attention deficit and hyperactivity disorder, autism and obsessive and compulsive disorder by modulating the dimer signaling to dMSNs.

## 5. Conclusion

This study presents for the first time evidence for an A_2A_R/H_3_R heterodimer based on functional complementation and co-immunoprecipitation assays in HEK-293 cells, where co-activation of the receptors leads to enhanced A_2A_R signaling and attenuation of H_3_R functionality. In rat striatal tissue the interaction occurs in synapses where H_3_R activation modifies the binding affinity of the A_2A_R.

## Acknowledgements

We thank Juan Escamilla-Sánchez and Raúl González-Pantoja for excellent technical assistance. R. M.-G. held a Conacyt graduate scholarship (244993).

## Author contributions

R. M.-G., R. v. R. and J.-A. A.-M. designed the study; R. M.-G., C. G.-R., J.-M. A. and M. T. R. performed the experiments; R. M.-G., J.-A. O.-R., R. v. R. and J.-A. A.-M. performed data analysis. R. M.-G., R. v. R. and J.-A. A.-M. wrote the manuscript. All authors revised and approved the manuscript.

## Legends for Supplementary Figures

**Supplementary Figure 1**. A. The activation of the Gα_s_-coupled histamine H_2_ receptor did not induce Ca^2+^ mobilization wen transfected alone, but did so when the Gα_qs_ was coexpressed, proving the functionality of the chimeric protein. B. The truncated H_3_Rs of 421 and 411 residues were unable to induce Ca^2+^ mobilization and functional complementation was not observed by homodimerization with either receptor. C. Functional complementation assay showing that the co-activation of the WT-H_3_R with increasing concentrations of RAMH and the A_2A_R_302_-Gα_qs_ with a fixed concentration of CGS-21680 did not modified the response. D. The co-activation of the H_3_R_427_-Gα_qi4_ and the WT-A_2A_R did not recover the Ca^2+^ response observed when the second construct was transfected alone.

**Supplementary Figure 2**. Effect of histamine and RAMH on the H_3_R and H_4_R and cAMP modulation of the A_2A_R/H_3_R heterodimer A. Histamine induced Ca^2+^ mobilization by activation of the H_3_R only when the Gα_qi4_ was expressed. B. Activation of the H_4_R with histamine caused a Ca^2+^ response promoted by the Gα_qi4_. No response was observed by the solely transfection of the H_4_R. C and D. Plots showing the cAMP formation experiments expressed as percentage of the basal. C. A_2A_R signaling is increased after co-activation with the H_3_R. No change in the CGS-21680 efficacy or potency was observed when H_3_R was co-expressed with A_2A_R. D. A_2A_R co-expression and co-activation caused a prevention of H_3_R-medited cAMP inhibition, suggesting the prevalence of A_2A_R signaling in the heterodimer. The means ± SEM are shown in table 2.

**Supplementary Figure 3**. Transmission electron microscopy of rat striatal synaptosomes. A. Abundant presence of isolated terminals (arrow heads). Other common components such as mitochondrion (m) and myelin fragments (M) can be observed. B. Close view of a presynaptic terminal, distinguishable by the synaptic vesicles (v) and mitochondria, making contact with a post-synaptic element (p). The pre-synaptic active zone and the post-synaptic density can be distinguished.

**Supplementary Figure 4**. Effect of the endogenous H_3_R agonist histamine on the A_2A_R affinity for its agonist [^3^H]CGS-21680. A. Histamine (1 μM) increased A_2A_R affinity for CGS-21680. Values are means ± SEM from 5 replicates from a representative experiment. B. Analysis of 4 independent experiments.The statistical analysis was performed with paired Student’s *t* test.

**Supplementary Table 1:**
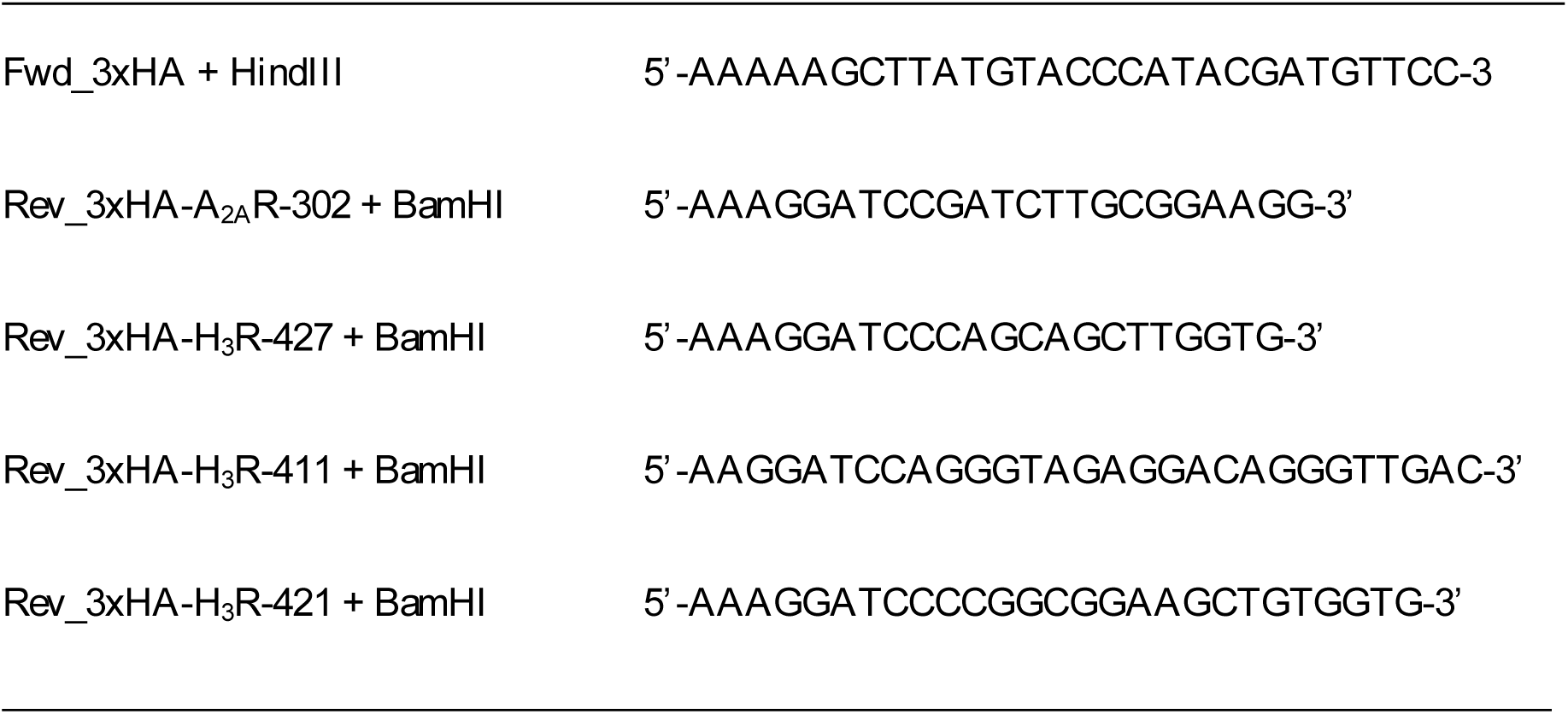
Primers employed in the generation of the truncated A_2A_ and H_3_ receptors

**Supplementary Table 2.** Pharmacological data of all the functional complementation assays performed.

| Transfection | Agonist | E<sub>max</sub> (%) | pEC<sub>50</sub> | EC<sub>50</sub> (nM) |
| --- | --- | --- | --- | --- |
| H<sub>3</sub>R | RAMH | ND | ND | ND |
|  | Histamine | ND | ND | ND |
| H<sub>3</sub>R + Gα<sub>qi4</sub> | RAMH | 419 ± 25 | 7.73 ± 0.31 | 19.95 |
|  | Histamine | 504 ± 22 | 6.19 ± 0.17 | 645.65 |
| A<sub>2A</sub>R | CGS | ND | ND | ND |
| A<sub>2A</sub>R + Gα<sub>qs4</sub> | CGS | ND | ND | ND |
| A<sub>2A</sub>R<sub>302</sub>-Gα<sub>qi4</sub> | CGS | ND | ND | ND |
| A<sub>2A</sub>R<sub>302</sub>-Gα<sub>qs4</sub> | CGS | ND | ND | ND |
| A<sub>2A</sub>R<sub>302</sub>-Gα<sub>qi4</sub> + A<sub>2A</sub>R | CGS | ND | ND | ND |
| A<sub>2A</sub>R<sub>302</sub>-Gα<sub>qs4</sub> + A<sub>2A</sub>R | CGS | ND | ND | ND |
| H<sub>3</sub>R<sub>427</sub>-Gα<sub>qi4</sub> | RAMH | 753 ± 41 | 7.14 ± 0.21 | 72.44 |
| H<sub>3</sub>R<sub>427</sub>-Gα<sub>qs4</sub> | RAMH | ND | ND | ND |
| H<sub>3</sub>R<sub>427</sub>-Gα<sub>qi4</sub> + H<sub>3</sub>R | RAMH | 523 ± 41 | 7.38 ± 0.34 | 41.68 |
| H<sub>3</sub>R<sub>427</sub>-Gα<sub>qs4</sub> + H<sub>3</sub>R | RAMH | ND | ND | ND |
| H<sub>3</sub>R<sub>411</sub>-Gα<sub>qi4</sub> | RAMH | ND | ND | ND |
| H<sub>3</sub>R<sub>421</sub>-Gα<sub>qi4</sub> | RAMH | ND | ND | ND |
| H<sub>3</sub>R<sub>421</sub>-Gα<sub>qi4</sub> + H<sub>3</sub>R | RAMH | ND | ND | ND |
| H<sub>411</sub>-Gα<sub>qi4</sub> + H<sub>3</sub>R | RAMH | ND | ND | ND |
| $A_{2A}R_{302}\text{-}G\alpha_{qi4} + H_3R$ | RAMH | $449 \pm 25$ | $7.31 \pm 0.23$ | 48.97 |
| | CGS | $429 \pm 14$ | $7.63 \pm 0.15$ | 23.44 |
|  | + RAMH |  |  |  |
| $A_{2A}R\text{-}G\alpha_{qs4} + H_3R$ | RAMH | ND | ND | ND |
| $H_3R_{427}\text{-}G\alpha_{qs4} + A_{2A}R$ | CGS | ND | ND | ND |
| $H_3R_{427}\text{-}G\alpha_{qi4} + A_{2A}R$ | CGS | ND | ND | ND |
|  | CGS | ND | ND | ND |
|  | + RAMH |  |  |  |
| $H_4R$ | Histamine | ND | ND | ND |
|  | RAMH | ND | ND | ND |
| $H_4R + G\alpha_{qi4}$ | Histamine | $725 \pm 51$ | $6.35 \pm 0.25$ | 446.68 |
| | RAMH | $551 \pm 27$ | $6.15 \pm 0.16$ | 794.32 |
| $A_{2A}R_{302}\text{-}G\alpha_{qi4} + H_3R$ | Histamine | ND | ND | ND |
| $A_{2A}R_{302}\text{-}G\alpha_{qi4} + H_4R$ | Histamine | ND | ND | ND |
| $A_{2A}R_{302}\text{-}G\alpha_{qi4} + D_2R$ | Quinpirole | $612 \pm 34$ | $6.47 \pm 0.22$ | 338.84 |
| $D_2R$ | Quinpirole | ND | ND | ND |
| $H_2R + G\alpha_{qs4}$ | Dimaprit | $860 \pm 37$ | $6.15 \pm 0.12$ | 707.94 |
| $H_2R$ | Dimaprit | ND | ND | ND |

Data are means ± SEM from 3 experiments. RAMH and CGS-21680 were tested at 100 nM. RAMH: R- $\alpha$ -methylhistamine; CGS: CGS-21680. ND; not determined.

